# Heat hypersensitivity of ryanodine receptor type 1 mutants implicated in malignant hyperthermia

**DOI:** 10.1101/2020.10.29.351452

**Authors:** Kotaro Oyama, Vadim Zeeb, Toshiko Yamazawa, Takashi Murayama, Hideto Oyamada, Yoshie Harada, Norio Fukuda, Shin’ichi Ishiwata, Madoka Suzuki

**Author notes:** K.O., V.Z., and T.Y. contributed equally to this work. **Corresponding Authors** Kotaro Oyama, Quantum Beam Science Research Directorate, National Institutes for Quantum and Radiological Science and Technology, 1233 Watanukimachi, Takasaki-shi, Gunma 370-1292, Japan, Vadim Zeeb, Institute of Theoretical and Experimental Biophysics, Russian Academy of Sciences, Pushchino, Moscow Region 142292, Russia, Madoka Suzuki, Institute for Protein Research, Osaka University 3-2 Yamadaoka, Suita, Osaka 565-0871, Japan.

## Abstract

Cellular heat-sensing is a universal strategy for avoiding thermal damage and adapting to environments by regulating thermogenic activities. If heat-sensing results in the acceleration of processes governing cellular thermogenesis, hyperthermia can occur. However, how this positive feedback loop contributes to hyperthermia development, especially the gap between heat-sensing and thermogenesis, remains largely unknown. Here, we show that an optically controlled local heat pulse induces an intracellular Ca^2+^ burst in cultured HEK 293 cells overexpressing ryanodine-receptor-type-1 (RyR1) mutants related to the life-threatening illness malignant hyperthermia (MH), and that the Ca^2+^ burst originates from heat-induced Ca^2+^-release (HICR) because of the mutant channels’ heat hypersensitivity. Furthermore, the heat hypersensitivity of the four RyR1 mutants was ranked, highlighting the complexity of MH. Our findings reveal the novel cellular heat-sensing mechanism, HICR, is essential for the functional positive feedback loop causing MH, suggesting a well-tuned HICR is fundamental for heat-mediated intracellular signaling.

## Introduction

Heat-sensing is essential for avoiding thermal tissue damage and regulating body temperature^1^. Molecular temperature sensors in mammals and insects are temperature-activated transient receptor potential (TRP) channels^2^. In plants, a reversible temperature-dependent liquid–liquid phase transition of intrinsically disordered proteins has recently been revealed as a thermosensing mechanism^3^. In addition to the absolute temperature, cells also sense how it changes over time. This sensing frequently appears as shifts in chemical and physical processes^4^ or as systems composed of cooperative or antagonistic protein activities. For instance, rapid heating excites cells due to heat-induced displacement of ions near the membranes^5,6^. Rapid cooling increases intracellular Ca^2+^ concentration ([Ca^2+^]_i_) in muscles and induces muscle contraction, a phenomenon known as rapid cooling contracture^7,8^. In cardiac muscles, heat increases both Ca^2+^-sensitivity and myofibril tension generation but decreases the amplitude of Ca^2+^ transients that result in hyperthermic negative inotropy^8,9^. Because of the many different types of heat-sensing mechanisms, one may expect a life-threatening illness when heat-sensing positively affects the processes governing cellular thermogenesis; the thermally accelerated heat release eventually elevates body temperature beyond normal, causing hyperthermia. However, no key molecule, process, or system has been identified as a heat sensor in diseases such as malignant hyperthermia (MH).

MH is known to be triggered by inhalation of anesthetic gases, depolarizing muscle relaxants, and rarely, exposure to high ambient temperature or vigorous exercise^10,11^. Typical symptoms of MH are body temperatures elevated above 39°C and increased muscle rigidity. These symptoms are fatal if not treated immediately.

MH is caused by mutations in ryanodine receptor type 1 (RyR1) Ca^2+^-release channels, dihydropyridine receptors, and Src-homologous-3 and cysteine-rich domain-containing protein 3 in skeletal muscles^10,12,13^. The majority of human MH-associated mutations have been identified in the RyR1 gene. Normal RyR1 releases Ca^2+^ from the sarcoplasmic reticulum (SR) during excitation–contraction coupling. In skeletal muscles expressing these mutants, enhanced Ca^2+^ release from the lumen of the SR elevates [Ca^2+^]_i_, and uncontrolled hypermetabolism and hyperthermia are induced^11^. Environmental heat stress also triggers MH in knock-in mice expressing RyR1 mutations^14–16^. Therefore, the sudden elevation of the body temperature strongly suggests mutual amplification between Ca^2+^ release and thermogenesis, i.e., a positive feedback loop, in the mechanism of MH development. However, the intrinsic feature of this positive feedback loop remains elusive because it is unclear how elevated body temperature affects Ca^2+^ release. In fact, experimental^17–20^ and computational^21^ analyses have demonstrated that elevated temperatures reduce Ca^2+^ release in wild type (WT) RyR1–3. These results contradict the situation in MH, in which Ca^2+^ release through mutant RyR1s is maintained even with an elevated body temperature. Furthermore, little is known about how MH onset occurs when the body temperature is still within the normal range and symptoms begin to appear.

Here, the Ca^2+^ release induced by heat is shown experimentally to be the key feature of the positive feedback loop between Ca^2+^ and thermogenesis. By applying optically controlled local heat pulses with a focused near-infrared (IR) laser beam^22–26^ to cultured human embryonic kidney (HEK) 293 cells overexpressing MH mutant RyR1s, we directly examined the heat sensitivities of the mutants. Ca^2+^ release through RyR1 is dependent on [Ca^2+^]_i_^27^. Therefore, with this single cell-based assay system using HEK 293 cells, one can examine the Ca^2+^ release through RyR1 mutants *in situ*, without myopathy development^28–41^. The expression levels of endogenous RyRs in HEK 293 cells is negligible compared to those of the expressed RyR1s^37,42^. Transient heating of the cells within the field of view can be carried out nearly simultaneously. Therefore, the following artefacts can be avoided: changes in cell morphology; drifting focus due to thermal expansion of components in the experimental set-up; photo-bleaching of fluorophores; relocation of Ca^2+^ indicators due to leakage from, or internalization by intracellular compartments; and thermal damage to biomolecules and cells caused by long periods of heat exposure. Moreover, the temperature changes can be controlled with sub-degree accuracy. We successfully compared the heat sensitivity of MH mutants quantitatively; the mutants displayed greater heat sensitivity than WT RyR1. Our results suggest that during MH onset, a small amount of heat stress can cause elevated Ca^2+^ release via the heat hypersensitive MH mutants, causing hypermetabolism and hyperthermia that accelerates Ca^2+^ release. Thus, the thermogenic cascade becomes unstoppable.

## Results

### Malignant hyperthermia mutants cause heat-pulse-induced Ca^2+^ release

HEK 293 cells expressing either WT or mutant RyR1s (Q156K, R164C, and Y523S) were selected across the rank order of the activity of RyR1 mutants with N-terminal region mutations—Q156K was amongst the lowest ranked, Y523S was among the highest ranked, and R164C had an intermediate ranking (WT<Q156K<R164C<Y523S)^35,37,40^—to test their capabilities as heat sensors (**Fig. 1a**). The rank order was introduced to compare the amount of Ca^2+^ leakage among RyR1 mutants^37^. A higher rank indicates higher [Ca^2+^]_i_ and thus lower [Ca^2+^] in the SR and endoplasmic reticulum (ER)^29,30,35,37^.

**Figure 1.**
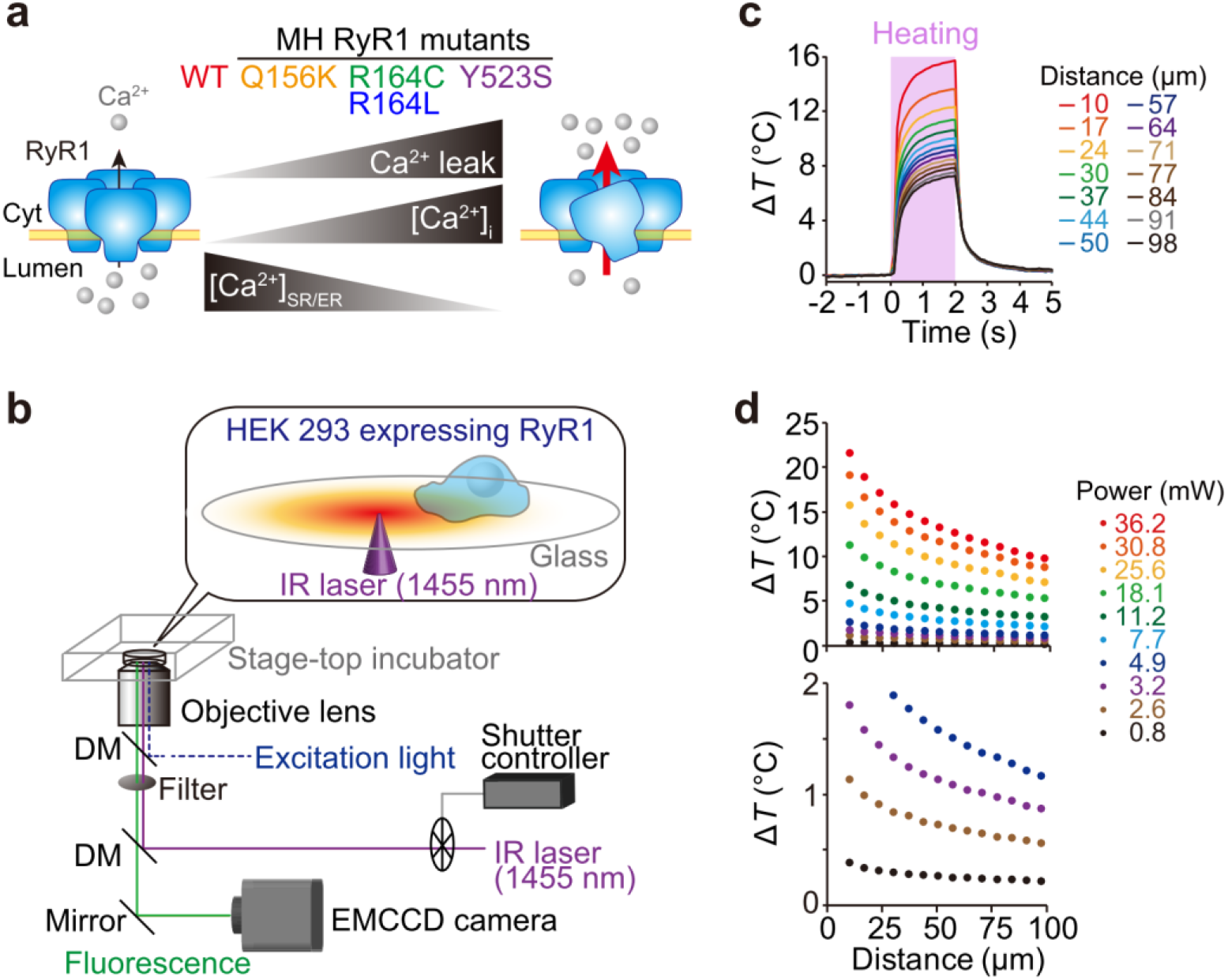
Experimental design to examine heat sensitivities of various malignant hyperthermia mutants of ryanodine receptor type 1. (**a**) Properties of malignant hyperthermia (MH) mutants of ryanodine receptor type 1 (RyR1) investigated in this study. Ca^2+^ leakage through mutant RyR1s (higher rank) is greater than that through wild-type (WT) receptors (the lowest rank). Cytoplasmic [Ca^2+^] ([Ca^2+^]_i_) of cells expressing RyR1 mutants is higher than that of cells expressing WT RyR1. The [Ca^2+^] in sarco/endoplasmic reticulum ([Ca^2+^]_SR/ER_) is depleted because of the Ca^2+^ leakage. The rank order was based on the literature^35,37^. (**b**) Schematic illustration of the fluorescence microscopy set-up. A 1455-nm infrared (IR) laser beam was guided to the sample stage by a dichroic mirror (DM) and objective lens and focused on the medium. The temperature in the field of view was elevated locally because of the absorption of IR light by the aqueous medium. Sample temperature was controlled by a stage-top incubator. (**c**) Time courses of temperature changes (Δ*T*) at various distances from the heat source. The pink region indicates the period of heat pulse. Laser power, 25.6 mW. (**d**) Temperature gradients formed by various laser powers. The bottom panel is the enlarged view of Δ*T*s between 0°C and 2°C.

Heat stimulation was applied using a 1455-nm IR laser^22–26,43^ while fluorescent Ca^2+^ imaging was performed (**Fig. 1b**). The temperature in the field of view was elevated almost immediately (<100 ms) after initiating heat stimulation, and it returned to its original level (re-cooling) when the stimulation was stopped after 2 seconds of heating (**Fig. 1c**). The amplitude Δ*T* was adjustable down to 1°C or less at the lower end by the laser’s power and distance from where it was focused (**Fig. 1d, Supplementary Fig. 1**); hence heat pulses of various Δ*T*s could be applied to cells in the same field of view (138 μm×138 μm) simultaneously.

No [Ca^2+^]_i_ changes were observed in control cells without induced RyR1 expression (−doxycycline (Dox)) or cells expressing WT RyR1 when heat pulses of Δ*T=*10±2°C were applied at the base temperature *T*_0_=24°C (**Fig. 2a**, **Supplementary Movie 1**). In contrast, there were rapid (<~500 ms), large [Ca^2+^]_i_ increases (Ca^2+^ bursts) after initiation of the heat pulse that continued after re-cooling in most R164C cells in the field of view (**Fig. 2b**, **Supplementary Movie 2**). In Q156K and Y523S cells, [Ca^2+^]_i_ decreased during the heat pulse, but Ca^2+^ bursts occurred relatively late (>~10 s) and soon (~2 s) after re-cooling, respectively **(Fig. 2c**, **Supplementary Fig. 2a**). The decrease in [Ca^2+^]_i_ during heating (apparent in WT cells in **Fig. 2a** and **Supplementary Movie 1**) could be attributable to heat-activated Ca^2+^ uptake by sarco/endoplasmic reticulum Ca^2+^-ATPase (SERCA)^23,44^. The maximum changes in the Ca^2+^ bursts, Δ*F*_max_/*F*_0_, during the 20 s after heating initiation were significantly larger in WT, Q156K, R164C, and Y523S cells than in −Dox cells (**Fig. 2d**). The experiments were repeated at physiological temperature, *T*_0_=36°C, with the same Δ*T* (10±1°C). The Ca^2+^ bursts observed in Q156K, R164C, and Y523S cells were similar to those observed at *T*_0_=24°C, whereas significant Ca^2+^ bursts were observed in −Dox and WT cells with mean Δ*F*_max_/*F*_0_ values comparable to that of Q156K (**Supplementary Fig. 2b**). To summarize, Δ*F*_max_/*F*_0_ and the fraction of cells exhibiting Ca^2+^ bursts were largely dependent on the RyR1 mutations and also on *T*_0_.

**Figure 2.**
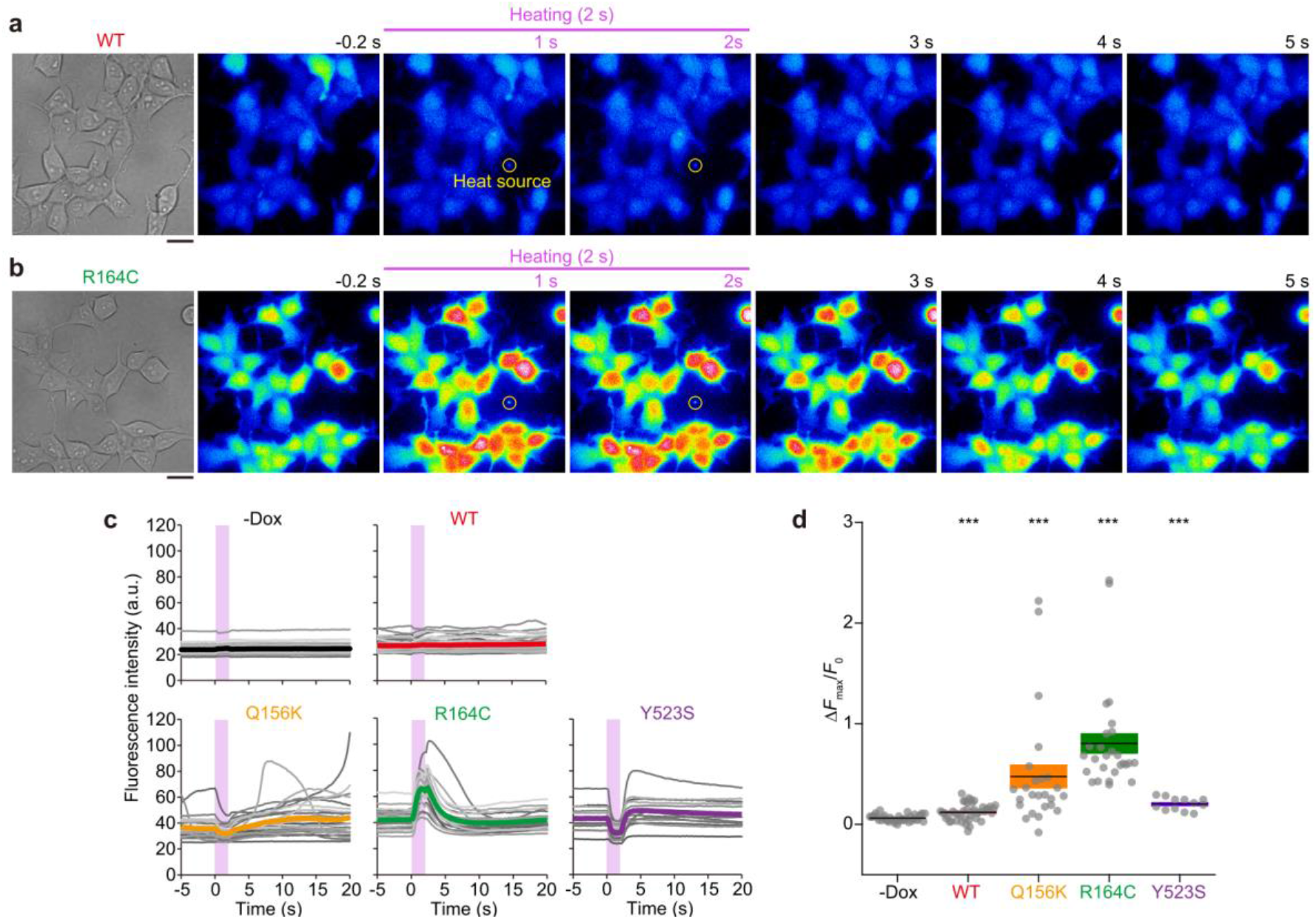
Intracellular Ca^2+^ responses to heat pulses in HEK 293 cells expressing ryanodine receptor type 1 mutants. (**a** and **b**) Bright-field and fluorescence images of fluo-4-loaded HEK 293 cells expressing WT RyR1 (**a**) and R164C (**b**). Background increase during heating was caused by IR laser beam scattering (apparent in **Supplementary Movie 1**). Yellow circles indicate the position of the heat source. Scale bars, 20 μm. (**c**) Changes in the fluorescence intensity of fluo-4. Each gray curve represents an individual cell. The thick colored curves indicate the average intensities. Pink regions indicate the periods of the heat pulses. (**d**) Maximum changes in relative fluorescence intensity of fluo-4 (Δ*F*_max_/*F*_0_) during the 20 s after heating initiation. Horizontal bars and error bars indicate the means±SEM. Statistical significance was determined by comparison with −Dox cells (*n*=40) using the Steel test (\*\*\**p*<0.001). WT, *n*=43 and *p*=3.5 × 10^−4^; Q156K, n=25 and *p*=5.3 × 10^−8^; R164C, n=27 and *p*=1.5 × 10^−11^; Y523S, n=12 and *p*=1.9 × 10^−6^. Laser power, 25.6 mW; Δ*T*=10±2°C; *T*_0_=24°C.

### Ca^2+^ flows from the endoplasmic reticulum to the cytosol primarily through ryanodine receptor type 1 during heat-induced Ca^2+^ bursts

The RyR1 mutation-dependent variation in heat-induced Ca^2+^ bursts suggests that the major Ca^2+^ source is the ER, and that Ca^2+^ flows from the lumen of ER to the cytosol through RyR1 channels. The major Ca^2+^ source was examined in WT RyR1, the mutant RyR1 in the middle of the rank order, R164C, and −Dox cells at physiological temperature, i.e., 36°C (**Fig. 3**). The Ca^2+^ burst was preserved in all WT, R164C and −Dox cells in Ca^2+^-free medium, whereas the burst was suppressed when Ca^2+^ was depleted from the ER by 2 μM thapsigargin (a SERCA inhibitor). These results support the idea that the Ca^2+^ source for the heat-induced Ca^2+^ burst is not the extracellular space but the ER.

**Figure 3.**
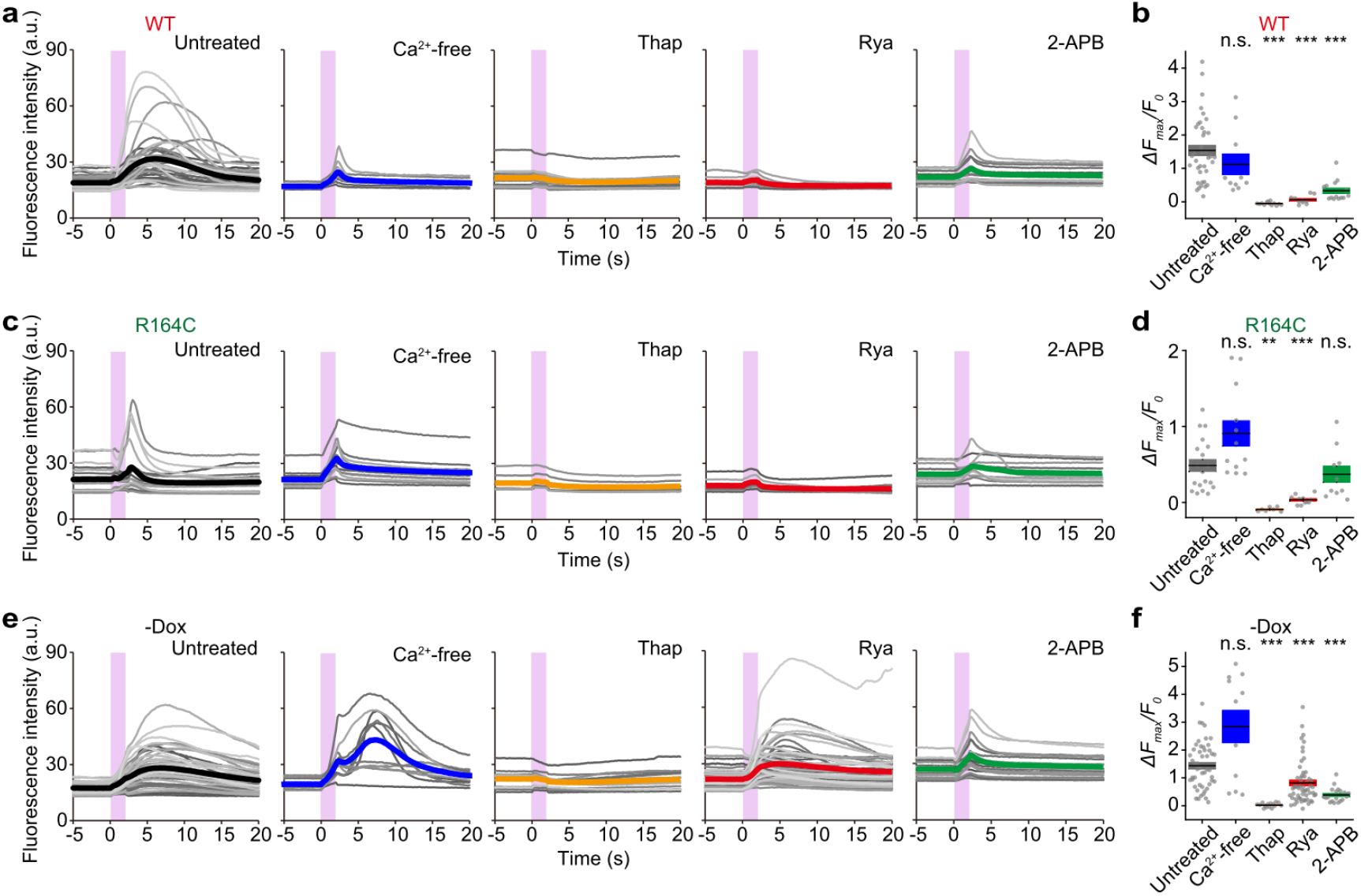
Contribution of ryanodine receptors and inositol trisphosphate receptors to heat-induced Ca^2+^ bursts. (**a, c,** and **e**) Time courses of the fluorescence intensity of fluo-4 in untreated cells, Ca^2+^-free solution (Ca^2+^-free), and in the presence of 2 μM thapsigargin (Thap), 100 μM ryanodine (Rya), or 100 μM 2-aminoethyl diphenylborinate (2-APB). Gray curves represent individual cells. The thick colored curves represent averages. Pink regions indicate the periods of the heat pulses. (a) WT, (c) R164C, (e) −Dox. (**b, d,** and **f**) Maximum changes in relative fluorescence intensity, Δ*F*_max_/*F*_0_. Horizontal bars and error bars indicate the means ± SEM. (b) WT, (d) R164C, (f) −Dox. Statistical significance was examined by comparison with the untreated cells using the Steel test (\*\**p*<0.01; \*\*\**p*<0.001; n.s., not significant). Data for WT, *p*=0.51 (Ca^2+^-free), 3.8 × 10^−7^ (Thap), 7.2 × 10^−6^ (Rya), and 1.4 × 10^−5^ (2-APB). Data for R164C, *p*=0.11 (Ca^2+^-free), 0.0013 (Thap), 9.7 × 10^−5^ (Rya), and 0.91 (2-APB). Data for −Dox, *p*=0.18 (Ca^2+^-free), 2.4 × 10^−8^ (Thap), 1.5 × 10^−5^ (Rya), and 2.3 × 10^−6^ (2-APB). Laser power, 25.6 mW; Δ*T*=10±1°C; *T*_0_=36°C.

The major Ca^2+^ channels on the ER membrane are RyRs and inositol trisphosphate receptors (IP_3_Rs). Ca^2+^ bursts in WT RyR1 and R164C cells were blocked by 100 μM ryanodine (inhibitor of RyR) (**Fig. 3a–d**), suggesting dominant contributions of respective RyR1 channels expressed in these cells. The suppression of Ca^2+^ bursts by ryanodine was much less effective in −Dox cells than in WT RyR1 and R164C cells (**Fig. 3e,f**); thus we confirmed that the contribution of the negligible endogenous RyR^45^ was relatively minor. The contribution of endogenous IP_3_R was also examined, as we have previously reported that IP_3_R plays a dominant role in heat-induced Ca^2+^ bursts after re-cooling in HeLa^44^ and WI-38^23^ cells. In fact, there was a significant suppression of Ca^2+^ bursts in −Dox cells by 100 μM 2-APB (unspecific inhibitor of IP_3_R^46–48^, of which the half maximal inhibition concentration (IC_50_) to IP_3_R is 42 μM^49^) (**Fig. 3e,f**), suggesting a major contribution of IP_3_R to Ca^2+^ bursts in −Dox cells. Although the peak intensity of fluo-4 (Δ*F*_max_/*F*_0_) was also reduced by 2-APB in WT RyR1 cells (**Fig. 3b)**, Ca^2+^ bursts were apparent (**Fig. 3a)**. Furthermore, no significant suppression of Ca^2+^ bursts was observed in R164C cells (**Fig. 3c,d)**. Therefore, we concluded that overexpressed RyR1 channels dominate in producing the Ca^2+^ bursts in their respective cells.

### Endoplasmic reticulum-targeted fluorescent Ca^2+^ probes indicate that the endoplasmic reticulum is a major source of released Ca^2+^

To further examine the hypothesis that the ER is the source of Ca^2+^ in the heat-induced Ca^2+^ burst, we monitored the [Ca^2+^] in the lumen of ER ([Ca^2+^]_ER_) using an ER-targeted fluorescent Ca^2+^ probe, G-CEPIA1*er*^50^, stably expressed in WT RyR1, Q156K, R164C, and Y523S cells. In WT RyR1, the fluorescence intensity of G-CEPIA1*er* decreased during the heat pulse (Δ*T*=10±1°C) and then recovered immediately after re-cooling (**Fig. 4a**, **Supplementary Fig. 3a, Supplementary Movie 3)**. The quick recovery after re-cooling suggests that the decrease during the heat pulse was because of thermal quenching of G-CEPIA1*er* fluorescence. In cells demonstrating relatively low G-CEPIA1*er* fluorescence intensity, the signal rise during heating was apparent because of IR-laser-beam scattering (e.g., R164C and Y523S in **Supplementary Fig. 3a**). A second decrease reached a minimum ~5 s after re-cooling and then recovered gradually. The initial value *F*_0_ of G-CEPIA1*er* and the second decrease were significantly reduced when Ca^2+^ was depleted from the ER lumen by 2 μM thapsigargin (**Fig. 4b**, **Supplementary Fig. 3a and b**). Furthermore, the second decrease appeared to coincide with the Ca^2+^ burst in the cytosol (**Supplementary Fig. 2a**). Hence, the second decrease of G-CEPIA1*er* fluorescence is considered to represent the [Ca^2+^]_ER_ decrease. The decrease in Q156K cells was comparable to that in WT cells, whereas those in R164C and Y523S cells were significantly lower (**Fig. 4b**). These results demonstrate positive correlations between the amplitudes of [Ca^2+^]_ER_ decrease (Δ*F*_min_/*F*_0_ of G-CEPIA1*er*) and that of the Ca^2+^ burst (Δ*F*_max_/*F*_0_ of fluo-4) (**Fig. 4c**), providing strong support for the conclusion that the heat-induced Ca^2+^ burst arises from Ca^2+^ release from the ER through RyR1.

**Figure 4.**
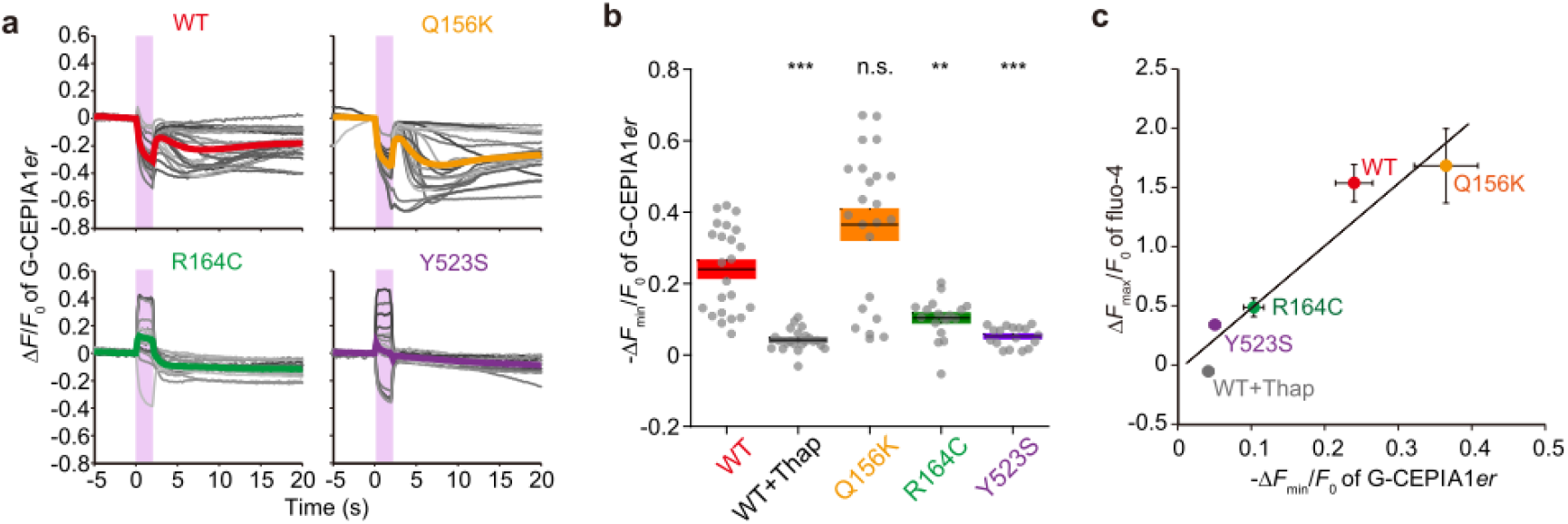
Heat-induced Ca^2+^ release from the endoplasmic reticulum. (**a**) Relative change in the fluorescence intensity, Δ*F*/*F*_0_, of G-CEPIA1*er* in cells expressing WT, Q156K, R164C, or Y523S. Data in **Supplementary Fig. 3a** were analyzed and plotted. *n*=23, 23, 18, and 16 cells for WT, Q156K, R164C, and Y523S, respectively. Pink regions indicate the periods of heat pulses. (**b**) Minimum relative change in G-CEPIA1*er* fluorescence intensity-Δ*F*_min_/*F*_0_ after heating. Statistical significance was determined by comparison with WT using the Steel test (\**p*<0.05; \*\**p*<0.01; \*\*\**p*<0.001). *p*=1.0 × 10^−7^ (WT + Thap), 0.086 (Q156K), 0.0035 (R164C), and 2.0 × 10^−6^ (Y523S). Horizontal bars and error bars indicate the means ± SEM. (**c**) Relationship between the change in [Ca^2+^]_er_ (−Δ*F*_min_/*F*_0_ of G-CEPIA1*er*) and [Ca^2+^]_i_ (Δ*F*_max_/*F*_0_ of fluo-4; **Supplementary Fig. 2**). The correlation coefficient *R*=0.96 (*p*=0.011). Laser power, 25.6 mW; Δ*T*=10±1°C; *T*_0_=36°C.

### Variation in heat sensitivity among wild-type and mutant ryanodine receptor type 1 channels and its ambient temperature sensitivity

Last, the heat sensitivities of WT and RyR1 mutants were examined systematically by exposing cells to heat pulses of variable Δ*T* and evaluating the fraction of cells exhibiting Ca^2+^ bursts (**Fig. 5a**, **Supplementary Fig. 4**). At *T*_0_=36°C, the Ca^2+^ burst was more frequently observed when Δ*T* was larger in WT and all mutant cells. However, there was a gradient among them. For example, approximately 40% of R164C cells showed Ca^2+^ bursts in response to a heat pulse of Δ*T*=1°C, but Ca^2+^ bursts were only observed in 20% or fewer WT and other mutant cells; i.e., R164C cells showed the highest heat sensitivity, Y523S and Q156K cells were less sensitive, and WT cells were the least sensitive. This variation in heat sensitivity was quantitatively compared using the threshold values of Δ*T* (Δ*T*_th_) at which 50% of the cells responded (**Fig. 5b**). This analysis is independent from spontaneous (non-thermal) [Ca^2+^]_i_ fluctuations and the amplitude of the Ca^2+^ burst, which can depend on [Ca^2+^]_ER_. Based on this data, one can draw a rank order in Δ*T*_th_ as R164C≪Y523S<Q156K<WT at both *T*_0_=24°C and 36°C (**Fig. 5b** **and** **c**, **Supplementary Figs 5 and 6a,b**). We further examined R164L, which has a rank-order of activity similar to that of R164C^37,40^, and found that the Δ*T*_th_ of R164L was comparable to that of R164C. It was also apparent that Q156K and WT cells were more heat-sensitive at *T*_0_=36°C than at *T*_0_=24°C.

**Figure 5.**
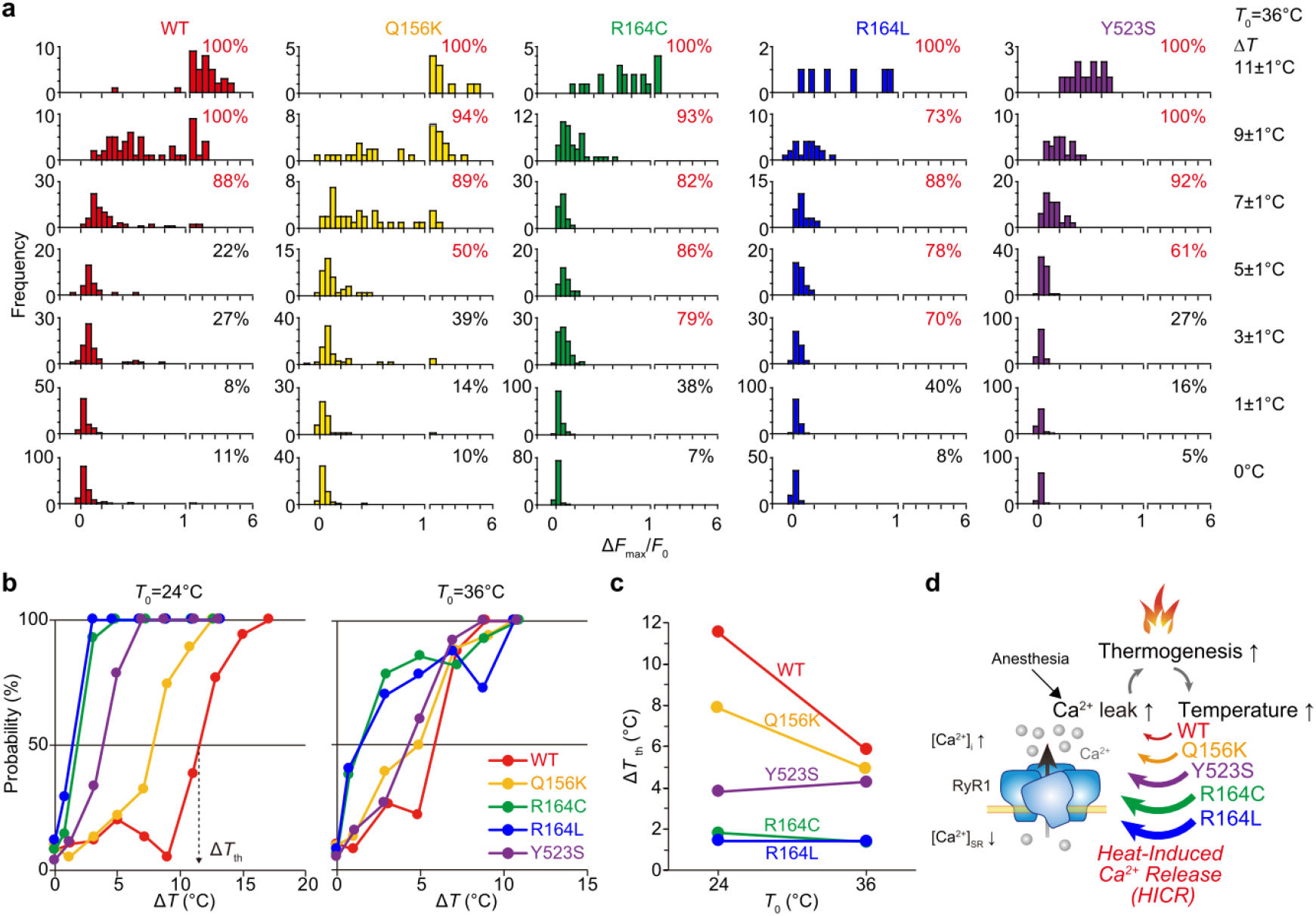
Rank order of ryanodine receptor type 1 mutants’ heat sensitivity. (**a**) Histograms showing [Ca^2+^]_i_ increases (Δ*F*_max_/*F*_0_s of fluo-4) in response to heat pulses of various amplitudes (Δ*T*). The numbers on the top right of each panel indicate the response probability that the cells showed significant [Ca^2+^]_i_ increase (Δ*F*_min_/*F*_0_>Δ*F*_th_). Data in **Supplementary Fig. 4** were analyzed and plotted. *T*_0_=36°C. (**b**) Relationship between Δ*T* and the response probability. The Δ*T*_th_ is defined as the Δ*T* at which the response probability=50%. (**c**) Relationship between *T*_0_ and Δ*T*_th_. Rank order in Δ*T*_th_ at *T*_0_=36°C was R164C (1.4°C)=R164L (1.4°C)<Y523S (4.3°C)<Q156K (4.9°C)<WT (5.8°C). The rank order at *T*_0_=24°C was R164L (1.4°C)≈R164C (1.8°C)<Y523S (3.8°C)<Q156K (7.9°C)<WT (11.5°C). (**d**) Schematic illustration of proposed positive-feedback loop closed by heat-induced Ca^2+^-release (HICR) explaining MH initiation and its aggravation. Anesthesia-triggered Ca^2+^ leak through MH RyR1 mutants induces hyperthermia. The temperature rise destabilizes RyR1 and causes HICR.

## Discussion

MH is a life-threatening illness caused by mutations in RyR1 that manifests as a sudden and unstoppable temperature elevation at the whole-body level. Here, we propose a link between the macroscopic whole-body temperature elevation of MH and the nanoscopic thermodynamics of RyR1 mutants. The current study successfully demonstrated heat-induced release of Ca^2+^ in selected MH RyR1 mutants using local heating with a focused IR laser beam, and that the mutant RyR1s are more heat-sensitive than WT RyR1s. The heat hypersensitivity and Ca^2+^ release induced by heat pulses in mutant RyR1 channels suggest that positive feedback between thermogenesis and Ca^2+^ release is responsible for MH progression. When Ca^2+^ leakage through MH RyR1 mutants is enhanced by MH inducers (e.g., inhalation of anesthetics), the resulting [Ca^2+^]_i_ increase leads to thermogenesis by SERCA^51^ and possibly also by Ca^2+^ leakage itself^52^. It is noteworthy that both SERCA and RyR1 are localized in the ER membrane. Then, the temperature elevation results in Ca^2+^ release through the channels, as demonstrated in the present study (for a schematic, see **Fig. 5d**). We propose naming this process heat-induced Ca^2+^-release (HICR). A critical role for HICR in the aggravation of MH is also suggested, especially at the initial stage, when low levels of heat stress originating from metabolism or exercise could initiate Ca^2+^ release because of the heat hypersensitivity of MH mutants. The HICR could coincide with the positive-feedback loop over a relatively longer time scale that Hamilton *et al*. suggested^53,54^. They showed that increased [Ca^2+^]_i_ activates the production of both reactive oxygen (ROS) and nitrogen (RNS) species in heterozygous knock-in mice with a Y524S (Y522S in human) mutation; these products then modify RyR1 and other skeletal muscle proteins to promote Ca^2+^ releases in an unclarified manner. Because of the relatively short time scale of HICR, the slower processes of ROS/RNS production and post-transcriptional modifications may contribute to the additional complexity of HICR and thus of MH. We found that Δ*T*_th_ is smaller in R164C and Y523S cells than in WT cells. This result is consistent with previous reports that muscle contraction and sudden death can be triggered by a moderate temperature increase in R165C (R163C in human) and Y524S knock-in mice that correspond to rabbit R164C and Y523S mutants, respectively^14,15^. Durham *et al.* have also reported that [Ca^2+^]_i_ was increased in the skeletal muscles of Y524S knock-in mice more than that in WT mice in response to the same degree of temperature elevation^53^. Among the mutants, the rank order did not necessarily reflect the order in Δ*T*_th_. The Y523S channel mutant is leakier than R164C(L) (**Fig. 1a**)^29,30,37^. However, the Δ*T*_th_ was greater in Y523S cells than in R164C(L) cells. According to previous studies examining caffeine-induced Ca^2+^ release^29,30,37^, one possible reason is that depletion of basal [Ca^2+^]_ER_ before the heat pulse reduces the amount of Ca^2+^ released from the ER. Because the Y523S mutant is the most destabilized channel, it would release more Ca^2+^, even without a heat pulse. Thus, the excess temperature elevation probably has a lesser effect on Ca^2+^ release through the Y523S channel than the R164C and R164L channels.

The heat hypersensitivity of MH mutants revealed in the present study could be caused by destabilized interactions between N-terminal domains (A, B and C) and neighboring domains at the “N-terminal hotspot” of residues 35–614, one of three known hotspots related to MH^55,56^ (**Supplementary Fig. 7**). The Q156 and R164 in domain A interact with domain B to form interface 1 within the ABC subunit at the N-terminus^57^. At the bottom of domain A (interface 4), R164 interacts with the core domain^57^. Y523 is exposed at the surface of domain C, facing the cytosolic shell domain (interface 3)^57^. One possible explanation for our data is that the inter-domain interactions are destabilized by heat in the mutants and cause Ca^2+^ leakage followed by intracellular Ca^2+^ bursts that are more heat-sensitive than in the WT. Hence, the heat hypersensitivity identified in the current study may be a common property of MH RyR1 mutants. This idea is supported by the literature because it has been reported that MH RyR1 mutant structures are generally unstable under environmental heat stress^57–59^. We have also reported that ER Ca^2+^ levels in R164C(L) and other N-terminal mutant cells were lower at 36°C than at room temperature relative to WT^37^. Further confirmation of this hypothesis would be a target for future experiments using molecular dynamics simulations, which have been useful for studying the effect of inter-domain interface mutations on the heat sensitivity of RyRs^58,60,61^.

In summary, we applied local heating and found that RyR1 heat-sensing results in HICR. Our results revealed that the heat hypersensitivity of MH mutants is dependent on the experimental temperature. Thus, a positive feedback loop can be formed between heat-sensing and thermogenesis to cause an abnormally elevated body temperature in MH. This hypothesis can explain the overlap between the MH mechanism and exertional heat stroke, in which hyperthermia is induced by physical exertion in patients expressing MH mutants^62–64^. Apart from the essential role of the anomalous positive feedback loop in hyperthermia progression, the current data also suggest that a well-tuned HICR could be fundamental to intracellular signaling.

## Materials and Methods

### Chemicals

Dulbecco’s Modified Eagle Medium (DMEM) (08488-55), L-glutamine (16948-04), and hygromycin (09287-84) were purchased from Nacalai Tesque Inc. (Kyoto, Japan). Fetal bovine serum (FBS) (10437-028), penicillin and streptomycin (15140-122), Lipofectamine 2000 (11668) and fluo-4 AM (F14217) were purchased from Thermo Fisher Scientific (Waltham, MA, USA). Blasticidin (ant-bl-1) was purchased from InvivoGen (San Diego, CA, USA). Collagen type 1 (IFP9660) was purchased from the Research Institute for the Functional Peptides (Yamagata, Japan). Doxycycline (Dox) (D9891), 2-aminoethoxydiphenyl borate (2-APB) (D9754) and poly(methyl methacrylate) (PMMA, M_w_~15,000) (200336) were purchased from Sigma-Aldrich (St. Louis, MO, USA). Thapsigargin (586005) was purchased from Merck (Darmstadt, Germany). Ryanodine (185-02821) was purchased from FUJIFILM Wako Pure Chemical Corporation (Osaka, Japan). Europium (III) thenoyltrifluoroacetonate trihydrate (Eu-TTA) (21392-96-1) was purchased from Acros Organics (Pittsburgh, PA, USA). The stocks of 1 mM fluo-4 AM in DMSO, 2 mM thapsigargin in DMSO, 10 mM ryanodine in distilled water, and 100 mM 2-APB in DMSO were stored at −20°C until used.

### Cell culture

HEK 293 cells stably transformed with rabbit skeletal muscle RyR1 or its mutants (Q156K, R164C/L, or Y523S; human Q155K, R163C/L, or Y522S) were generated as reported previously^35,37,65^. The expression of RyR1 is inducible by Dox using the Flp-In T-Rex system (Thermo Fisher Scientific); hence the system is suitable for investigating the functions of fatal RyR1 mutants in living cells. The expression levels of WT and mutant RyR1s were similar.

The HEK 293 cells were cultured in flasks or on dishes coated with collagen (TPP Techno Plastic Products AG, Trasadingen, Switzerland) in culture medium (DMEM containing 10% FBS, 2 mM L-glutamine, 100 units ml^−1^ penicillin, and 100 μg ml^−1^ streptomycin) with added 100 μg ml^−1^ hygromycin and 15 μg ml^−1^ blasticidin at 37°C in 5% CO_2_. To coat the flasks and dishes with collagen, they were filled with a 0.001% collagen type 1 solution in distilled water for 1 h at 37°C. The flasks and dishes were washed with the fresh culture medium just before culturing the cells.

### Ca^2+^ imaging

Cells were seeded on collagen-coated glass base dishes (AGC Techno Glass, Shizuoka, Japan) at 37°C in 5% CO_2_ for 1–3 days. To induce RyR1 expression, the culture medium was replaced with the medium containing 2 μg ml^−1^ Dox. Cells were incubated for 24–36 h before the experiments were initiated.

Cytoplasmic Ca^2+^ dynamics were studied using the fluorescent Ca^2+^ probe fluo-4 AM. The cells were incubated in HEPES–buffered saline (HBS) (140 mM NaCl, 5 mM KCl, 1 mM MgCl_2_, 1 mM Na_2_HPO_4_, 10 mM HEPES, 2 mM CaCl_2_, 5 mM D-glucose, pH 7.4, adjusted with NaOH) containing 1 μM fluo-4 AM for 30 min at room temperature. The solution was replaced with fresh HBS, and the cells were incubated under the microscope for at least 10 min before observation to allow temperature stabilization to 24±1°C or 36±0.5°C. Where indicated, the cells were observed in Ca^2+^-free solution (140 mM NaCl, 5 mM KCl, 1 mM MgCl_2_, 1 mM Na_2_HPO_4_, 10 mM HEPES, 5 mM D-glucose, 2 mM EGTA, pH 7.4, adjusted with NaOH) at least 15 min after incubation. To inhibit the activity of SERCA or IP_3_R, the cells were incubated in HBS containing 1 μM fluo-4 AM and either 2 μM thapsigargin or 100 μM 2-APB for 30 min at 24°C and then observed in HBS containing 2 μM thapsigargin or 100 μM 2-APB. To inhibit RyR1, the fluo-4-loaded cells were observed in HBS containing 100 μM ryanodine.

To monitor Ca^2+^ dynamics in the ER, HEK 293 cells stably expressing G-CEPIA1*er*^50^ were prepared as follows. Lentiviral vectors harboring the G-CEPIA1*er* construct were produced by replacing the cDNA of enhanced green fluorescent protein (GFP) in pCL20c-MSCV-AcGFP-WPRE (kindly provided by Dr. Y. Ohashi, The University of Tokyo, Tokyo, Japan)^66^. HEK 293T cells were co-transfected with four plasmids, pCAGkGP1.1R, pCAG4RTR2, pCAG-VSV-G, and pCL20c-MSCV-G-CEPIA1er-WPRE, using Lipofectamine 2000. The cells were incubated at 37°C in 5% CO_2_ for 16 h. Then the culture medium was replaced, and the cells were incubated at 37°C in 5% CO_2_ for 36 h. The lentivirus-containing medium was collected and cleared by centrifugation at 1,500 rpm for 5 min at 4°C. To concentrate the lentivirus, the supernatant was centrifuged at 10,000 rpm at 4°C overnight. The pellets were suspended in 50 μl phosphate buffered saline and stored at −80°C until used. HEK 293 cells expressing WT RyR1 or the mutant RyR1s were transfected with the virus for G-CEPIA1*er*. Approximately 80%–95% of the cells were transduced. After at least two additional passages, the cells were used in the experiments.

### Optical set-up

The microscope with local heating apparatus was set up as described in our previous work^22–26,43^. Briefly, the local temperature around the cell was increased by a 1455-nm IR laser beam that is efficiently absorbed by water (KPS-STD-BT-RFL-1455-02-CO; Keopsys, Lannion, France). The duration of irradiation was controlled by a mechanical shutter (SSH-C4B; Sigma Koki, Tokyo, Japan). The laser power was measured using a thermal disk sensor (LM-3; Coherent, Santa Clara, CA, USA) and a power meter (FieldMaster; Coherent) at the level of the sample after passage through the objective lens. The permeability of the IR laser beam, which is the ratio of the output laser power at the level of the sample to the input laser power, was approximately 1.6%. Fluo-4, G-CEPIA1*er*, and Eu-TTA were excited by a solid-state illuminator (SPECTRA Light Engine; Lumencor, Beaverton, OR, USA; 377/50 nm for Eu-TTA and 485/20 nm for fluo-4 and G-CEPIA1*er*). The fluorescence and the bright-field images were observed with an inverted microscope (IX70; Olympus, Tokyo, Japan), equipped with a dichroic mirror (DM505; Olympus), an emission filter (BA515IF; Olympus), an objective lens (PlanApo N 60×/1.45 Oil, Olympus), and an electron multiplying charge coupled device (EMCCD) camera (iXon^EM^ + 897; Andor Technology, Belfast, UK). The temperature of the solutions on the sample stage was adjusted to 36±0.5°C using a thermostatically controlled incubator (INUCP-KRi-H2-F1; Tokai Hit, Shizuoka, Japan); otherwise the temperature of the solutions was 24±1°C.

### Analyses

The microscopic images were analyzed with ImageJ (National Institutes of Health, Bethesda, MD, USA).

The changes in local temperature were measured by thermal quenching of Eu-TTA that had been spin-coated on a glass base dish by a solution containing 5 mg ml^−1^ Eu-TTA and 10 mg ml^−1^ PMMA in acetone^23–25^. Relative changes in the intensity of Eu-TTA were calculated by Δ*F*/*F*_0_=(*F*_heating_−*I*_laser_)/(*F*_before−_*I*_back_)−1, where *F*_heating_ was the intensity at the end of the heating period, i.e., just before the IR laser beam was shut off, and *I*_laser_ was the background intensity caused by light scattering of the IR laser beam. *I*_back_ was the background intensity when the excitation light was off. Photo-bleaching was corrected by fitting a single exponential curve. Δ*F*/*F*_0_ of Eu-TTA was converted to Δ*T* using the relationship between Δ*T* and Δ*F*/*F*_0_ (−2.7% °C^−1^ at 24°C and −4.1% °C^−1^ at 36°C)^23,25^.

To calculate changes in the fluorescence intensities of fluo-4 and G-CEPIA1*er*, the outlines of cells were manually tracked in the bright-field images. Then the fluorescence intensities of fluo-4 and G-CEPIA1*er* were measured within the outlined areas. The distance between the area center and a laser spot was defined as the distance between the cell and the heat source. The change in the fluorescence intensity (Δ*F*) of fluo-4 was calculated from *F*−*F*_before_, where *F* was the fluorescence intensity at an arbitrary time, and *F*_before_ was the intensity just before heating was initiated (i.e., 10 s after beginning the observation). The basal fluorescence intensity of fluo-4 (*F*_0_) was calculated from *F*_before_−*I*_back_. The peak intensity of fluo-4 (Δ*F*_max_/*F*_0_) was calculated from the maximum Δ*F*/*F*_0_ during the 20 s after heating initiation. Light scattering by the IR laser beam (**Supplementary Movie 1**) was subtracted to calculate Δ*F*_max_/*F*_0_ during heating. No noticeable photo-bleaching of fluo-4 was observed during measurement (**Fig. 2a**).

The Δ*F*_max_/*F*_0_ of spontaneous [Ca^2+^]_i_ fluctuations in **Supplementary Fig. 8** was calculated by (*F*_max_−*F*_before_)/(*F*_before_−*I*_back_). To equalize the exposure time of excitation light (485 nm) and that in the heating experiments, *F*_before_ was set to the fluorescence intensity 10 s after starting the observation. *F*_max_ was the maximum intensity obtained from 10 to 30 s after starting the observation. The cumulative probability of Δ*F*_max_/*F*_0_ was fitted by a cumulative distribution function of the Gaussian distribution^67^

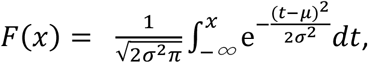

where μ and σ are the mean and the standard deviation (SD) of the Gaussian distribution, respectively. The fitting by least squares methods was performed in Excel 2016 (Microsoft, Redmond, WA, USA) using the following equation:

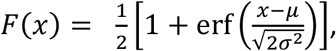

where erf(*x*) is an error function. The threshold Δ*F*_th_ of Δ*F*_max_/*F*_0_ was defined as μ+1.96σ. If Δ*F*_max_/*F*_0_ induced by a heat pulse was higher than Δ*F*_th_, the [Ca^2+^]_i_ increase response induced by the heat pulse was considered significant.

The threshold of Δ*T* (Δ*T*_th_) was defined as the Δ*T* that induced significant [Ca^2+^]_i_ increases (Δ*F*_max_/*F*_0_>Δ*F*_th_) in 50% of cells (**Fig. 5b**).

The Δ*F*_min_/*F*_0_ of G-CEPIA1*er* was calculated from (*F*_min_–*F*_before_)/(*F*_before_–*I*_back_), where *F*_min_ was the minimum intensity of G-CEPIA1*er* obtained from 2.4 s to 10 s after heating initiation. In this calculation of Δ*F*_min_/*F*_0_, photo-bleaching was corrected by fitting with a single exponential curve in Excel 2016 (Microsoft).

### Statistical analysis

Multiple groups were compared using the Steel test. For comparisons of two independent samples, the Mann–Whitney *U* test was used. These tests were performed using EZR (ver 1.51)^68^. Statistical significance was described by *p* values. Linear regression analysis was performed using OriginPro2016 software (OriginLab, Northampton, MA, USA).

## Acknowledgments

We sincerely thank the late Professor David H. MacLennan (University of Toronto, Toronto, ON, Canada) for valuable suggestions about useful mutagenesis strategies for *RyR1* cDNA constructs and constant encouragement. We also thank Ms. Tomomi Arai (Waseda University) for technical assistance. This work was supported by JSPS KAKENHI Grant Numbers 25707035 (to K.O.), 19K07306 (to T.Y.), JP22227005 (to S.I.), and 19H03198 (to M.S.), by the Russian Advanced Research Foundation, Grant Volkhov-A (to V.Z.), by the Japan Science and Technology Agency JPMJPR17P3 (to K.O.) and JPMJPR15F5 (to M.S.), and by the Human Frontier Science Program RGP0047/2018 (to V.Z. and M.S.).

## Author contributions

K.O., V.Z., T.Y., S.I., and M.S. designed the research study; K.O., V.Z., and T.Y. performed the experiments; T.M., H.O., J.S., K.K., and M.I. contributed new reagents/analytic tools;

K.O. analyzed the data; K.O., V.Z., T.Y., Y.H., N.F., S.I., and M.S. contributed to data analysis, and all authors were involved in drafting this work.

## Competing interests

The authors declare no conflicts of interest.

## Notes

### Competing Interest Statement

The authors have declared no competing interest.

## References

1. Tan, C. L. & Knight, Z. A. Regulation of body temperature by the nervous system. Neuron 98, 31–48 (2018).

2. Patapoutian, A., Peier, A. M., Story, G. M. & Viswanath, V. ThermoTRP channels and beyond: mechanisms of temperature sensation. Nat. Rev. Neurosci. 4, 529–539 (2003).

3. Jung, J.-H. et al. A prion-like domain in ELF3 functions as a thermosensor in Arabidopsis. Nature 585, 256–260 (2020).

4. Kondepudi, D. & Prigogine, I. Modern thermodynamics: from heat engines to dissipative structures. (John Wiley & Sons, Hoboken, 1998).

5. Shapiro, M. G., Homma, K., Villarreal, S., Richter, C. P. & Bezanilla, F. Infrared light excites cells by changing their electrical capacitance. Nat. Commun. 3, 736 (2012).

6. Liu, Q., Frerck, M. J., Holman, H. A., Jorgensen, E. M. & Rabbitt, R. D. Exciting cell membranes with a blustering heat shock. Biophys. J. 106, 1570–1577 (2014).

7. Sakai, T. Rapid cooling contracture. Jpn. J. Physiol. 36, 423–431 (1986).

8. Bers, D. M.. Excitation-contraction coupling and cardiac contractile force. (Kluwer Academic Publishers, Dordrecht, 2001).

9. Ishii, S. et al. Thermal activation of thin filaments in striated muscle. Front. Physiol. 11, 278 (2020).

10. Lanner, J. T., Georgiou, D. K., Joshi, A. D. & Hamilton, S. L. Ryanodine receptors: structure, expression, molecular details, and function in calcium release. Cold Spring Harb. Perspect. Biol. 2, a003996 (2010).

11. Rosenberg, H., Pollock, N., Schiemann, A., Bulger, T. & Stowell, K. Malignant hyperthermia: a review. Orphanet J. Rare Dis. 10, 93 (2015).

12. Riazi, S., Kraeva, N. & Hopkins, P. M. Malignant hyperthermia in the post-genomics era: new perspectives on an old concept. Anesthesiology 128, 168–180 (2018).

13. Ibarra Moreno, C. A. et al. An assessment of penetrance and clinical expression of malignant hyperthermia in individuals carrying diagnostic ryanodine receptor 1 gene mutations. Anesthesiology 131, 983–991 (2019).

14. Yang, T. et al. Pharmacologic and functional characterization of malignant hyperthermia in the R163C RyR1 knock-in mouse. Anesthesiology 105, 1164–1175 (2006).

15. Chelu, M. G. et al. Heat- and anesthesia-induced malignant hyperthermia in an RyR1 knock-in mouse. FASEB J. 20, 329–330 (2006).

16. Lopez, J. R., Kaura, V., Diggle, C. P., Hopkins, P. M. & Allen, P. D. Malignant hyperthermia, environmental heat stress, and intracellular calcium dysregulation in a mouse model expressing the p.G2435R variant of RYR1. Br. J. Anaesth. 121, 953–961 (2018).

17. Sitsapesan, B. Y. R. et al. Sheep cardiac sarcoplasmic reticulum calcium-release channels: modification of conductance and gating by temperature. J. Physiol. 434, 469–488 (1991).

18. Ferrier, G. R., Smith, R. H. & Howlett, S. E. Calcium sparks in mouse ventricular myocytes at physiological temperature. Am. J. Physiol. Heart Circ. Physiol. 285, H1495–H1505 (2003).

19. Protasi, F. et al. All three ryanodine receptor isoforms generate rapid cooling responses in muscle cells. Am J Physiol Cell Physiol 286, C662–C670 (2004).

20. Fu, Y. et al. Temperature dependence and thermodynamic properties of Ca^2+^ sparks in rat cardiomyocytes. Biophys. J. 89, 2533–2541 (2005).

21. Moskvin, A. S. S., Iaparov, B. I. I., Ryvkin, A. M. M., Solovyova, O. E. E. & Markhasin, V. S. Electron-conformational transformations govern the temperature dependence of the cardiac ryanodine receptor gating. JETP Lett. 102, 62–68 (2015).

22. Oyama, K. et al. Microscopic heat pulses induce contraction of cardiomyocytes without calcium transients. Biochem. Biophys. Res. Commun. 417, 607–612 (2012).

23. Itoh, H., Oyama, K., Suzuki, M. & Ishiwata, S. Microscopic heat pulse-induced calcium dynamics in single WI-38 fibroblasts. BIOPHYSICS 10, 109–119 (2014).

24. Shintani, S. A., Oyama, K., Fukuda, N. & Ishiwata, S. High-frequency sarcomeric auto-oscillations induced by heating in living neonatal cardiomyocytes of the rat. Biochem. Biophys. Res. Commun. 457, 165–170 (2015).

25. Oyama, K. et al. Triggering of high-speed neurite outgrowth using an optical microheater. Sci. Rep. 5 16611 (2015).

26. Oyama, K. et al. Directional bleb formation in spherical cells under temperature gradient. Biophys. J. 109, 355–364 (2015).

27. Endo, M. Calcium-induced calcium release in skeletal muscle. Physiol. Rev. 89, 1153–1176 (2009).

28. Tong, J. et al. Caffeine and halothane sensitivity of intracellular Ca^2+^ release is altered by 15 calcium release channel (ryanodine receptor) mutations associated with malignant hyperthermia and/or central core disease. J. Biol. Chem. 272, 26332–26339 (1997).

29. Tong, J., McCarthy, T. V.& MacLennan, D. H. Measurement of resting cytosolic Ca^2+^ concentrations and Ca^2+^ store size in HEK-293 cells transfected with malignant hyperthermia or central core disease mutant Ca^2+^ release channels. J. Biol. Chem. 274, 693–702 (1999).

30. Avila, G. & Dirksen, R. T. Functional effects of central core disease mutations in the cytoplasmic region of the skeletal muscle ryanodine receptor. J. Gen. Physiol. 118, 277–290 (2001).

31. Brini, M. et al. Ca^2+^ signaling in HEK-293 and skeletal muscle cells expressing recombinant ryanodine receptors harboring malignant hyperthermia and central core disease mutations. J. Biol. Chem. 280, 15380–15389 (2005).

32. Jiang, D. et al. Reduced threshold for luminal Ca^2+^ activation of RyR1 underlies a causal mechanism of porcine malignant hyperthermia. J. Biol. Chem. 283, 20813–20820 (2008).

33. Sato, K., Pollock, N. & Stowell, K. M. Functional studies of *RYR1* mutations in the skeletal muscle ryanodine receptor using human *RYR1* complementary DNA. Anesthesiology 112, 1350–1354 (2010).

34. Sato, K., Roesl, C., Pollock, N. & Stowell, K. M. Skeletal muscle ryanodine receptor mutations associated with malignant hyperthermia showed enhanced intensity and sensitivity to triggering drugs when expressed in human embryonic kidney cells. Anesthesiology 119, 111–118 (2013).

35. Nakano, M. et al. Construction and expression of ryanodine receptor mutants relevant to malignant hyperthermia patients in Japan. Showa Univ. J. Med. Sci. 26, 27–38 (2014).

36. Gomez, A. C. & Yamaguchi, N. Two regions of the ryanodine receptor calcium channel are involved in Ca^2+^-dependent inactivation. Biochemistry 53, 1373–1379 (2014).

37. Murayama, T. et al. Divergent activity profiles of type 1 ryanodine receptor channels carrying malignant hyperthermia and central core disease mutations in the amino-terminal region. PLoS One 10, e0130606 (2015).

38. Murayama, T. et al. Genotype-phenotype correlations of malignant hyperthermia and central core disease mutations in the central region of the RYR1 channel. Hum. Mutat. 37, 1231–1241 (2016).

39. Gomez, A. C., Holford, T. W. & Yamaguchi, N. Malignant hyperthermia-associated mutations in the S2-S3 cytoplasmic loop of type 1 ryanodine receptor calcium channel impair calcium-dependent inactivation. Am. J. Physiol. - Cell Ph. 311, C749–C757 (2016).

40. Yamazawa, T. et al. Insights into channel modulation mechanism of RYR1 mutants using Ca^2+^ imaging and molecular dynamics. J. Gen. Physiol. 152, e201812235 (2020).

41. Iyer, K. A. et al. Structural mechanism of two gain-of-function cardiac and skeletal RyR mutations at an equivalent site by cryo-EM. Sci. Adv. 6, eabb2964 (2020).

42. Oyama, K. et al. Single-cell temperature mapping with fluorescent thermometer nanosheets. J. Gen. Physiol. 152, e201912469 (2020).

43. Ishii, S. et al. Microscopic heat pulses activate cardiac thin filaments. J. Gen. Physiol. 151, 860–869 (2019).

44. Tseeb, V., Suzuki, M., Oyama, K., Iwai, K. & Ishiwata, S. Highly thermosensitive Ca^2+^ dynamics in a HeLa cell through IP3 receptors. HFSP J. 3, 117–123 (2009).

45. Nakamura, N., Yamazawa, T., Okubo, Y. & Iino, M. Temporal switching and cell-to-cell variability in Ca^2+^ release activity in mammalian cells. Mol. Syst. Biol. 5, 247 (2009).

46. Bootman, M. D. et al. 2-Aminoethoxydiphenyl borate (2-APB) is a reliable blocker of store-operated Ca^2+^ entry but an inconsistent inhibitor of InsP3-induced Ca^2+^ release. FASEB J. 16, 1145–1150 (2002).

47. Bilmen, J. G., Wootton, L. L., Godfrey, R. E., Smart, O. S. & Michelangeli, F. Inhibition of SERCA Ca^2+^ pumps by 2-aminoethoxydiphenyl borate (2-APB): 2-APB reduces both Ca^2+^ binding and phosphoryl transfer from ATP, by interfering with the pathway leading to the Ca^2+^-binding sites. Eur. J. Biochem. 269, 3678–3687 (2002).

48. Hu, H. Z. et al. 2-Aminoethoxydiphenyl borate is a common activator of TRPV1, TRPV2, and TRPV3. J. Biol. Chem. 279, 35741–35748 (2004).

49. Maruyama, T., Kanaji, T., Nakade, S., Kanno, T. & Mikoshiba, K. 2APB, 2-aminoethoxydiphenyl borate, a membrane-penetrable modulator of Ins(l,4,5)P3-induced Ca^2+^ release. J. Biochem. 122, 498–505 (1997).

50. Suzuki, J. et al. Imaging intraorganellar Ca^2+^ at subcellular resolution using CEPIA. Nat. Commun. 5, 4153 (2014).

51. de Meis, L., Arruda, A. P. & Carvalho, D. P. Role of sarco/endoplasmic reticulum Ca^2+^-ATPase in thermogenesis. Biosci. Rep. 25, 181–190 (2005).

52. Tullis, A., Block, B. A. & Sidell, B. D. Activities of key metabolic enzymes in the heater organs of scombroid fishes. J. Exp. Biol. 161, 383–403 (1991).

53. Durham, W. J. et al. RyR1 S-nitrosylation underlies environmental heat stroke and sudden death in Y522S RyR1 knockin mice. Cell 133, 53–65 (2008).

54. Lanner, J. T. et al. AICAR prevents heat-induced sudden death in RyR1 mutant mice independent of AMPK activation. Nat. Med. 18, 244–251 (2012).

55. Treves, S., Jungbluth, H., Muntoni, F. & Zorzato, F. Congenital muscle disorders with cores: the ryanodine receptor calcium channel paradigm. Curr. Opin. Pharmacol. 8, 319–326 (2008).

56. Hwang, J. H., Zorzato, F., Clarke, N. F. & Treves, S. Mapping domains and mutations on the skeletal muscle ryanodine receptor channel. Trends Mol. Med. 18, 644–657 (2012).

57. Tung, C.-C., Lobo, P. A., Kimlicka, L. & Van Petegem, F. The amino-terminal disease hotspot of ryanodine receptors forms a cytoplasmic vestibule. Nature 468, 585–588 (2010).

58. Kimlicka, L., Lau, K., Tung, C.-C. & Van Petegem, F. Disease mutations in the ryanodine receptor N-terminal region couple to a mobile intersubunit interface. Nat. Commun. 4, 1506 (2013).

59. Lau, K. & Van Petegem, F. Crystal structures of wild type and disease mutant forms of the ryanodine receptor SPRY2 domain. Nat. Commun. 5, 5397 (2014).

60. Zheng, W. & Liu, Z. Investigating the inter-subunit/subdomain interactions and motions relevant to disease mutations in the N-terminal domain of ryanodine receptors by molecular dynamics simulation. Proteins 85, 1633–1644 (2017).

61. Zheng, W. & Wen, H. Investigating dual Ca^2+^ modulation of the ryanodine receptor 1 by molecular dynamics simulation. Proteins (2020). doi:10.1002/prot.25971

62. Poussel, M. et al. Exertional heat stroke and susceptibility to malignant hyperthermia in an athlete: evidence for a link? J. Athl. Train. 50, 1212–1214 (2015).

63. Roux-Buisson, N. et al. Identification of variants of the ryanodine receptor type 1 in patients with exertional heat stroke and positive response to the malignant hyperthermia in vitro contracture test. Br. J. Anaesth. 116, 566–568 (2016).

64. Laitano, O., Murray, K. O. & Leon, L. R. Overlapping mechanisms of exertional heat stroke and malignant hyperthermia: evidence vs. conjecture. Sport. Med. 50, 1581–1592 (2020).

65. Murayama, T. et al. Role of amino-terminal half of the S4-S5 linker in type 1 ryanodine receptor (RyR1) channel gating. J. Biol. Chem. 286, 35571–35577 (2011).

66. Ohashi, Y. et al. A bicistronic lentiviral vector-based method for differential transsynaptic tracing of neural circuits. Mol. Cell. Neurosci. 46, 136–147 (2011).

67. Kobirumaki-Shimozawa, F. et al. Nano-imaging of the beating mouse heart in vivo: Importance of sarcomere dynamics, as opposed to sarcomere length per se, in the regulation of cardiac function. J. Gen. Physiol. 147, 53–62 (2016).

68. Kanda, Y. Investigation of the freely available easy-to-use software ‘EZR’ for medical statistics. Bone Marrow Transplant. 48, 452–458 (2013).

