## Supplemental Information for "Heat hypersensitivity of ryanodine receptor type 1 mutants implicated in malignant hyperthermia"

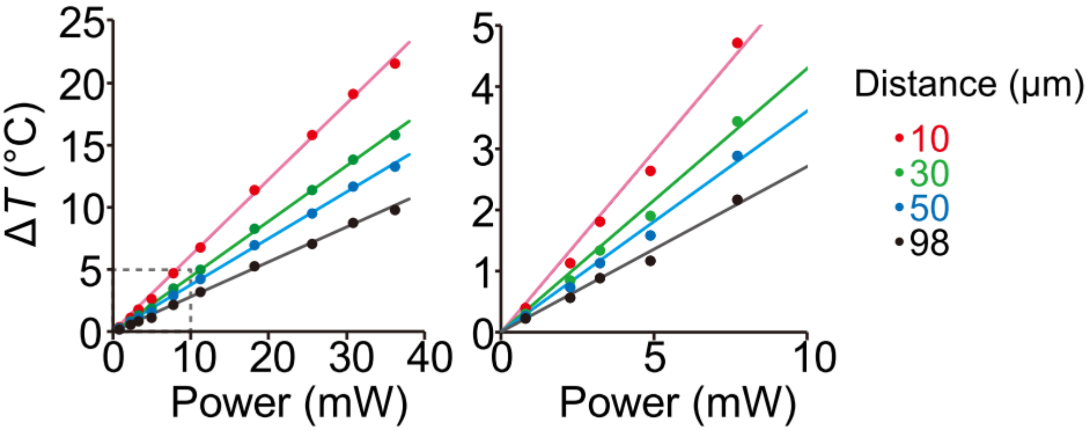


**Supplementary Figure 1. Relationship between laser power and temperature elevation (Δ*T*) at various distances from the heat source.** The Δ*T* was measured on the surface of a glass base dish by the thermal quenching of the temperature-sensitive dye Eu-TTA. Right, enlarged view of the area demarcated by dashed lines in the left panel. Error bars representing standard errors (SEM) (*n*=3 measurements) overlap the symbols.

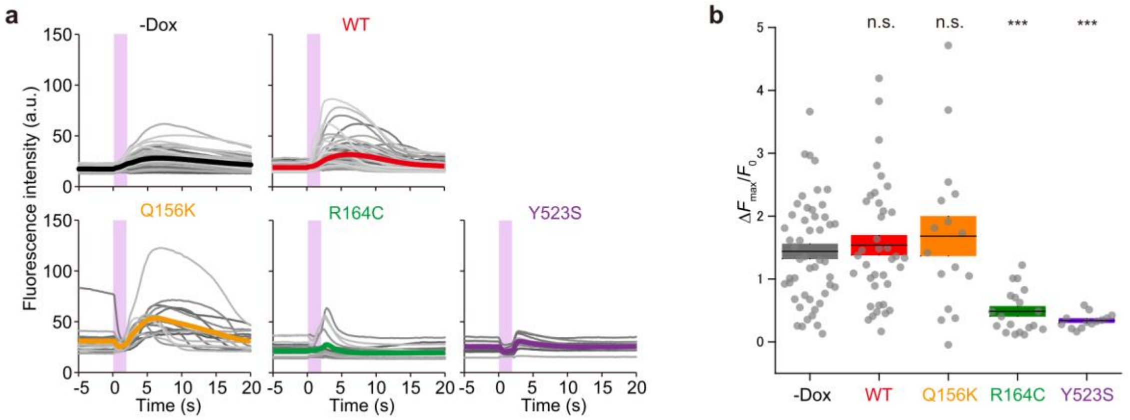


**Supplementary Figure 2. Intracellular $Ca^{2+}$ responses to a heat pulse in HEK 293 cells expressing ryanodine receptor type 1 mutants at physiological temperature.** (**a**) Time courses of the changes in fluorescence intensity of fluo-4 in HEK 293 cells with or without (−Dox) induced RyR1 expression at 36°C. Pink regions indicate the periods of the heat pulses. (**b**) The maximum changes in relative fluorescence intensity, $\Delta F_{max}/F_0$, were analyzed from data in (**a**). Horizontal bars and error bars indicate the means±SD. Statistical significance was determined by comparison with −Dox cells ($n$=49) using the Steel test (***$p$<0.001; n.s., not significant). WT, $n$=38 and $p$=0.99; Q156K, $n$=16 and $p$=0.97; R164C, $n$=19 and $p=1.9 \times 10^{-5}$; Y523S, $n$=13 and $p=1.9 \times 10^{-5}$. Laser power, 25.6 mW; $\Delta T$=10±1°C; $T_0$=36°C.

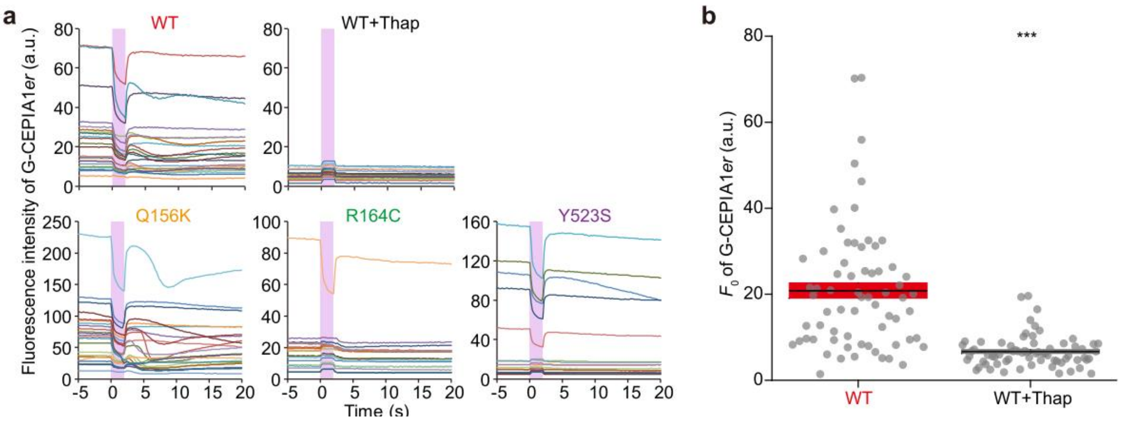


**Supplementary Figure 3. $Ca^{2+}$ dynamics in the endoplasmic reticulum of individual cells.** (**a**) Time courses of the fluorescence intensity of G-CEPIA1*er* in HEK 293 cells expressing WT RyR1 with or without 2 μM thapsigargin and in cells expressing RyR1 mutants. Pink regions indicate the periods of the heat pulses. Each line represents an individual cell. (**b**) G-CEPIA1*er* fluorescence intensity without heating, $F_0$. For untreated and thapsigargin-treated cells, $n$=67 and 69 cells, respectively. Statistical significance was examined using the Mann–Whitney $U$ test (***$p$<0.001). $p=4.7 \times 10^{-11}$. Laser power, 25.6 mW; $\Delta T$=10±1°C; $T_0$=36°C.

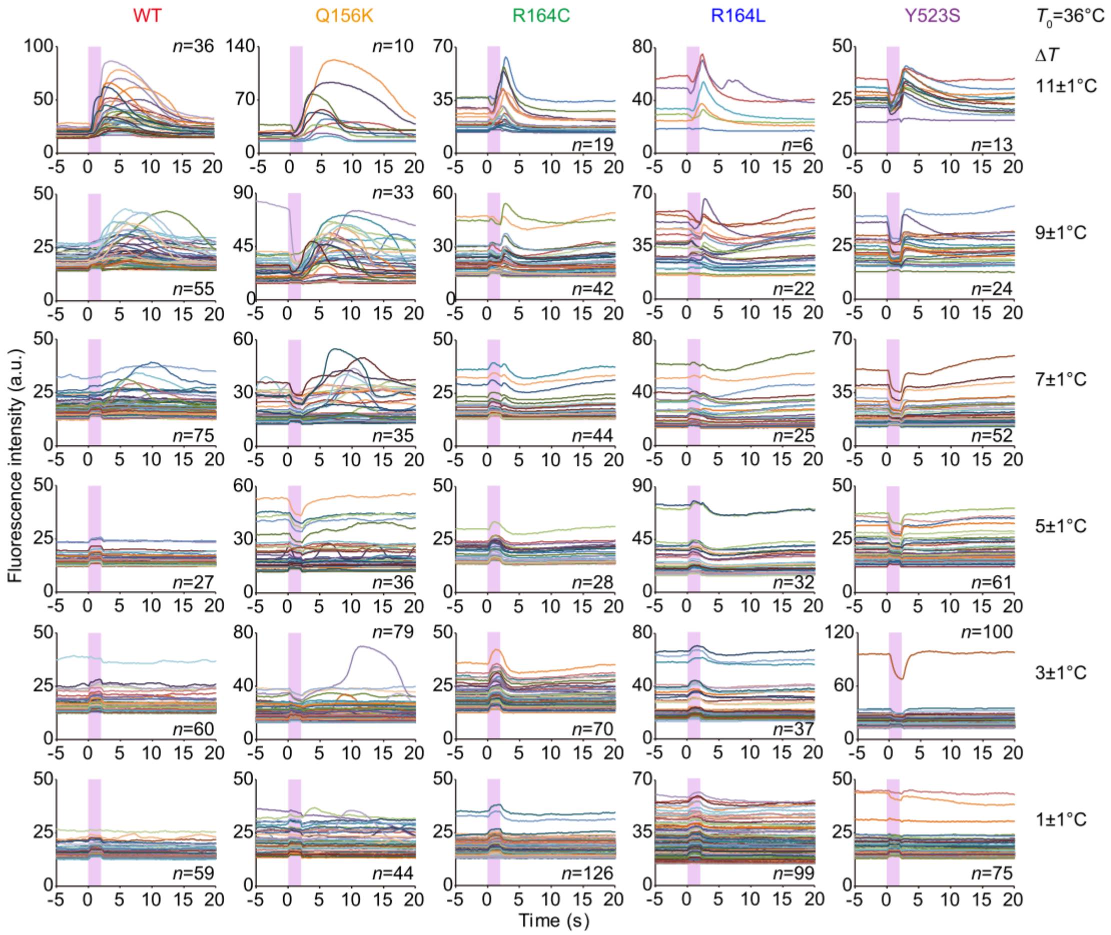


**Supplementary Figure 4. Intracellular $Ca^{2+}$ responses to heat pulses of various amplitudes in individual cells at physiological temperature.** Time courses of fluorescence intensity of fluo-4 in HEK 293 cells expressing WT RyR1 (left column) or RyR1 mutants at $T_0$=36°C. Pink regions indicate the periods of heat pulses.

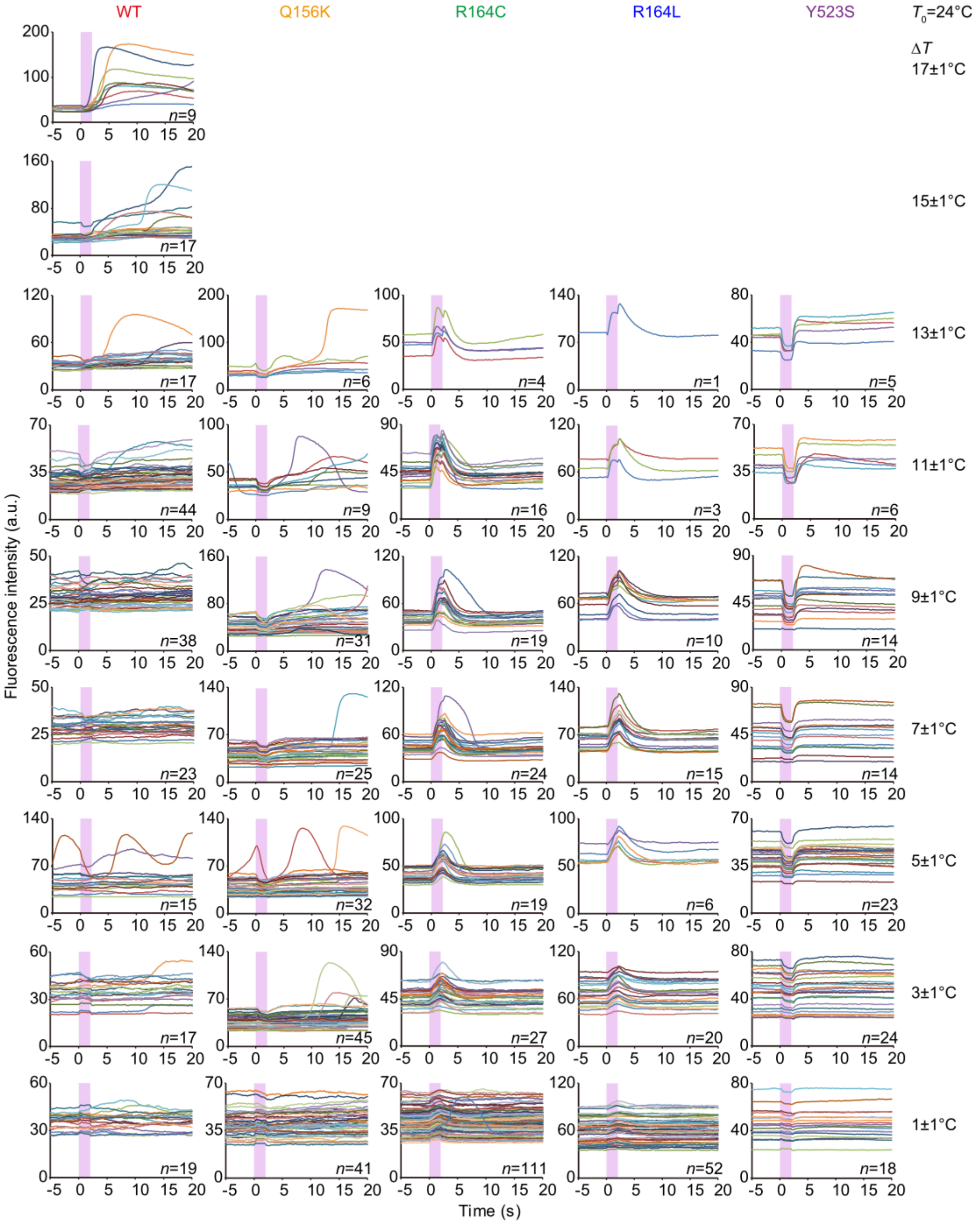


**Supplementary Figure 5. Intracellular $Ca^{2+}$ responses to heat pulses of various amplitudes in individual cells at room temperature.** Time courses of the fluorescence intensity of fluo-4 in HEK 293 cells expressing WT RyR1 (left column) or RyR1 mutants at $T_0$=24°C. Pink regions indicate the periods of the heat pulses.

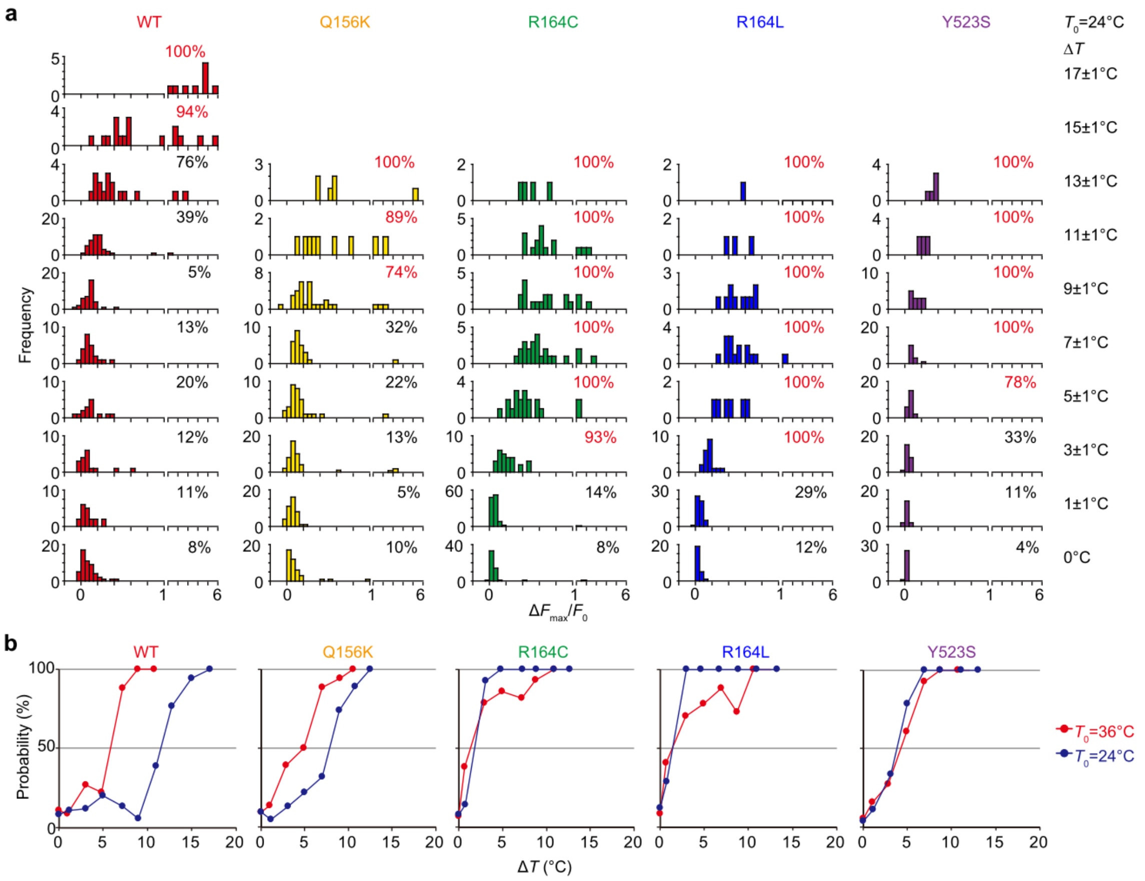


**Supplementary Figure 6. Intracellular $Ca^{2+}$ responses to heat pulses of various amplitudes.** (**a**) Histograms showing $[Ca^{2+}]_i$ increases ($\Delta F_{max}/F_0$ of fluo-4) in response to heat pulses of various amplitudes ($\Delta T$). The numbers in the top right of each panel indicate the response probability that the cells showed significant $[Ca^{2+}]_i$ increases ($\Delta F_{min}/F_0 > \Delta F_{th}$). Data in **Supplementary Fig. 5** were analyzed and plotted. $T_0$=24°C. (**b**) The relationship between $\Delta T$ and the response probability in various cell types at $T_0$=24°C and 36°C.

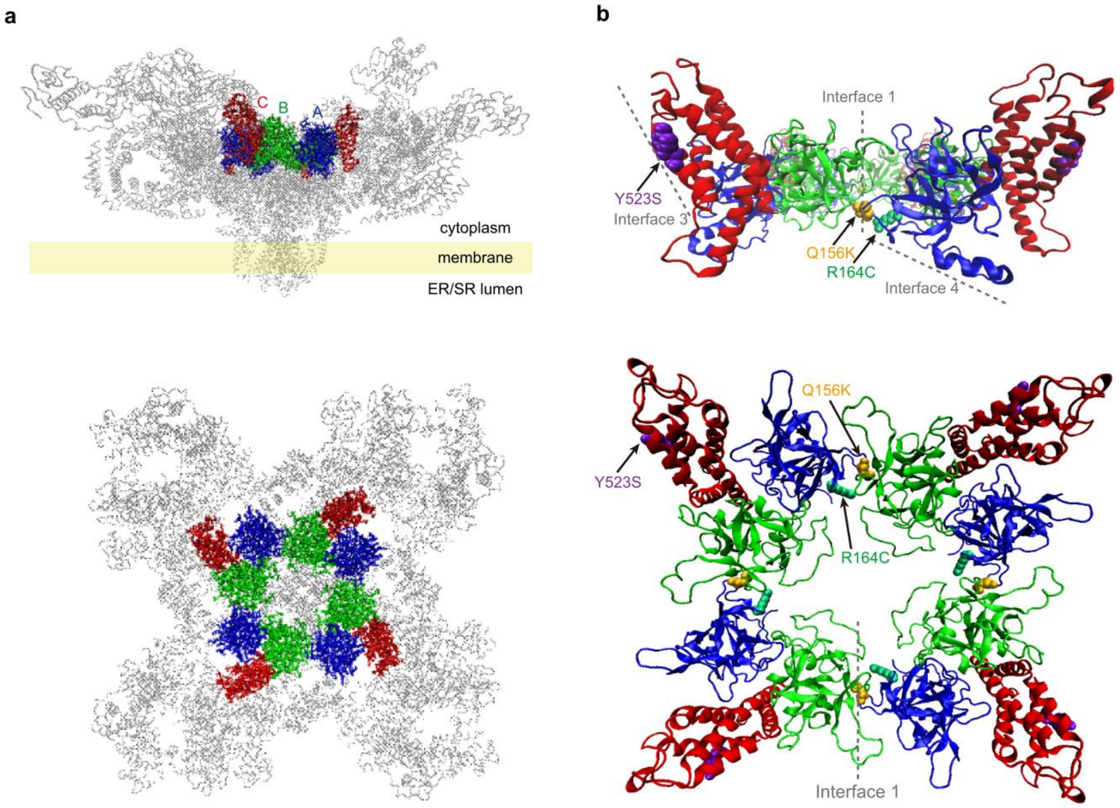


**Supplementary Figure 7. Structure of ryanodine receptor type 1.** (**a**) Overall view of RyR1 (PDB ID: 5GKZ)[1] structure. Side view (top) and the top view seen from the cytoplasm (bottom). The crystal structure of the N-terminal domain (1–559, NTD) tetramer (PDB ID: 2XOA)[2] is colored. The NTD subdomains (A, B, and C) are shown in blue, green, and red, respectively. (**b**) Enlarged view of the NTD tetramer. Side view (top) and the top view seen from the cytoplasm side (bottom). The mutation sites (R164, Q156, and Y523) and the interacting interfaces[2] are labeled. Interfaces 2, 5, and 6 are not shown because the mutations of the current study (R164, Q156, and Y523) are not located in these interfaces.

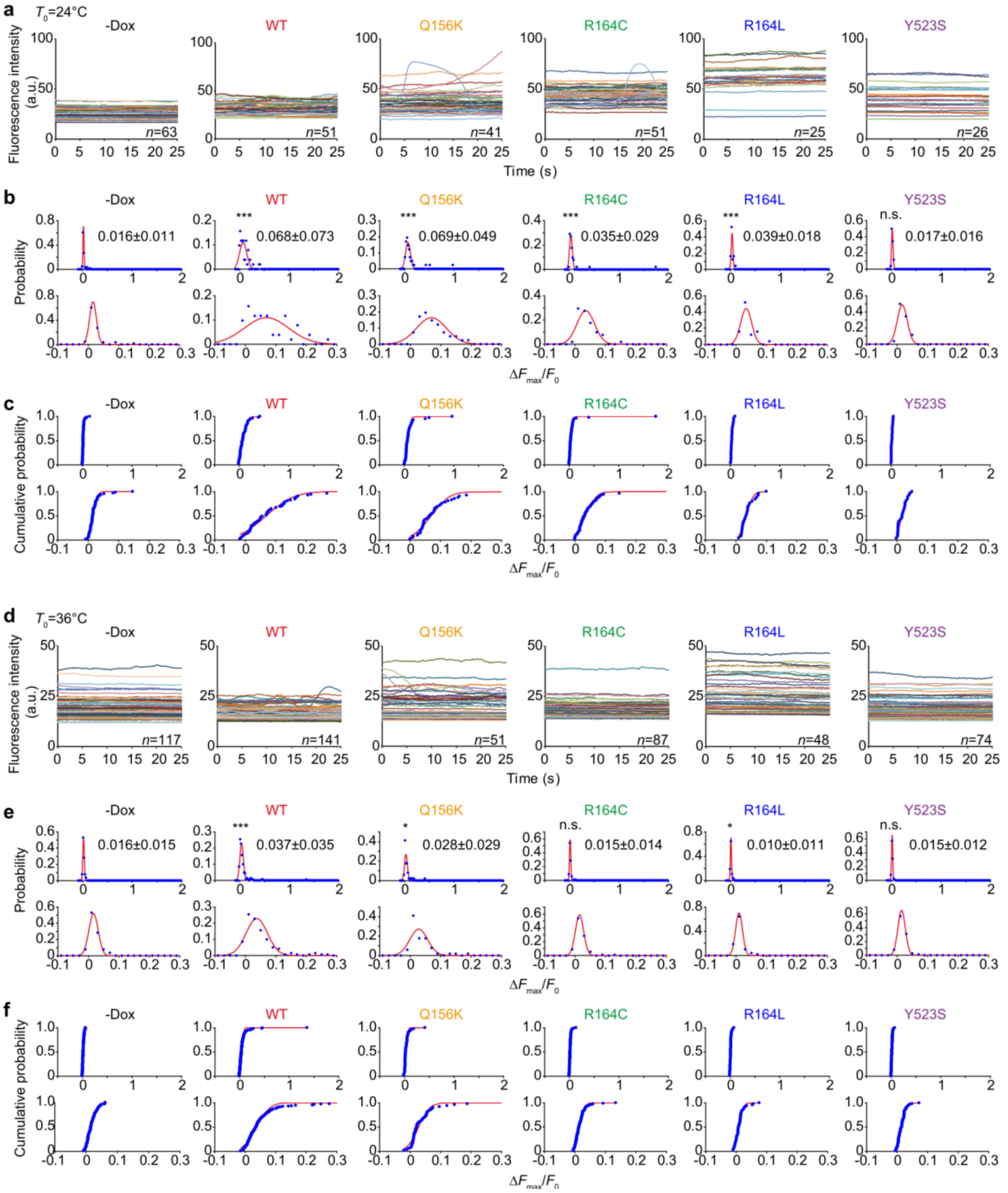


**Supplementary Figure 8. Endogenous fluctuations of $[Ca^{2+}]_i$ in HEK 293 cells expressing ryanodine receptor type 1 mutants.** (**a**) Time courses of changes in relative fluorescence intensity $\Delta F/F_0$ of fluo-4 in HEK 293 cells with or without induced expression of RyR1. $T_0$=24°C. (**b** and **c**) Histograms (**b**) and the cumulative histograms (**c**) of the maximum changes in relative fluorescence intensity ($\Delta F_{max}/F_0$) of fluo-4. Data in (**a**) were analyzed and plotted. Data (blue plots) were fitted by a single Gaussian function (**b**) or a cumulative distribution function of the Gaussian distribution (**c**) (red lines). See Materials and Methods for details of the fitting procedure. Numbers in (**b**) indicate the means ± SD, which were used to determine $\Delta F_{th}$ (see Materials and Methods). (**d**) Same as (**a**) but $T_0$=36°C. (**e** and **f**) Same as (**b**) and (**c**), respectively, but the data in (**d**) were analyzed and plotted. In (**b**) and (**e**), statistical significance was examined with −Dox cells using the Steel test (*$p$<0.05; ***$p$<0.001; n.s., not significant). In (**b**), $p$=3.0 × $10^{-6}$ (WT), 6.4 × $10^{-11}$ (Q156K), 2.3 × $10^{-4}$ (R164C), 3.0 × $10^{-5}$ (R164L), and 1.00 (Y523S). In (**e**), $p$=2.8 × $10^{-7}$ (WT), 0.0027 (Q156K), 0.81 (R164C), 0.0032 (R164L), and 0.87 (Y523S).

**Supplementary References**

1. des Georges, A. *et al.* Structural basis for gating and activation of RyR1. *Cell* **167**, 145–157.e17 (2016).
2. Tung, C. C., Lobo, P. A., Kimlicka, L. & Van Petegem, F. The amino-terminal disease hotspot of ryanodine receptors forms a cytoplasmic vestibule. *Nature* **468**, 585–588 (2010).

**Legends of Supplementary Movies**

**Movie 1. $Ca^{2+}$ response to a heat pulse in HEK 293 cells expressing wild-type ryanodine receptor type 1.** Fluorescence microscope images of fluo-4-loaded cells expressing wild-type ryanodine receptor type 1. Scale bar, 20 µm; laser power, 25.6 mW; $T_0$=24°C.

**Movie 2. Heat-induced $Ca^{2+}$ bursts in HEK 293 cells expressing R164C.** Fluorescence microscope images of fluo-4-loaded HEK 293 cells expressing R164C. Scale bar, 20 µm; laser power, 25.6 mW; $T_0$=24°C.

**Movie 3. Heat-induced endoplasmic reticulum $Ca^{2+}$ dynamics in HEK 293 cells expressing wild-type ryanodine receptor type 1.** Fluorescence microscope images of G-CEPIA1*er* in HEK 293 cells expressing wild-type ryanodine receptor type 1. Scale bar, 20 µm; laser power, 25.6 mW; $T_0$=36°C.
